# Effective Markovian dynamics method of solving non-Markovian dynamics of stochastic gene expression

**DOI:** 10.1101/2024.12.30.630775

**Authors:** Youming Li, Chen Jia

## Abstract

Experiments have shown that over 10% of proteins are degraded non-exponentially. Gene expression models for non-exponentially degraded proteins are notoriously difficult to solve since the underlying stochastic dynamics is non-Markovian. Here we develop an effective Markovian dynamics (EMD) method which converts a large class of non-Markovian models into effective Markovian ones so that they have the same mRNA and protein distributions at any fixed time. Using the EMD approach, we analytically solve some classical gene expression models with non-exponential or delayed protein decay, whose exact distributions are previously unknown and fail to be obtained using conventional queueing theory. Our theory successfully explains why non-exponentially degraded proteins on average have smaller mRNA-protein correlation than exponentially degraded proteins, and it predicts that bimodality is significantly enhanced in the presence of delayed protein degradation.

## Introduction

In living cells, protein degradation is vital for maintaining cellular homeostasis by eliminating unnecessary or damaged protein molecules. Proteins can be degraded by either the proteasome or lysosome, with proteins arriving at the lysosome either via autophagy or endocytosis [1]. All these degradation pathways share significant interdependence and rely on the availability of free ubiquitin. The ubiquitin-proteasome system (UPS) is responsible for *>* 80% of intracellular protein degradation in eukaryotes [2]. Recent studies have shown that *>* 10% of proteins in mammalian cells display non-exponential degradation, with the UPS system being involved in the degradation process for the majority of non-exponentially degraded (NED) proteins [3, 4].

Protein degradation often occurs through a sequence of events that are mediated by a complex proteolytic pathway, and hence many previous studies assume a time delay in this process [5–8]. Gene expression systems with delayed degradation are also widely used to model the dynamics of nascent RNA [9–11] since the conversion time of nascent RNA to mature RNA has been shown to be fairly deterministic [12]. However, gene expression models with non-exponential or delayed degradation are difficult to solve since the underlying stochastic dynamics is non-Markovian. The traditional method of solving such models is based on queueing theory [11]. Using queueing theory, analytical results have only been obtained in some simple models with renewal arrivals for protein synthesis and deterministic delay for protein degradation. However, thus far, there is still a lack of analytical results for some classical gene expression models such as the two-stage model with transcription and translation, as well as the three-stage model with transcription and translation, and gene state switching [13].

In this study, we propose a novel method of solving a large class of stochastic gene expression models with non-exponential or delayed protein degradation. The method is then applied to obtained analytical results for some important models that fail to be solved using queueing theory. Some testable predictions are also made.

### Two-stage model

Let *G* denote the gene, let *M* denote the mRNA, and let *P* denote the protein. We first focus on a two-stage stochastic gene expression model with non-exponential protein decay (Fig. 1(a)). The reaction scheme is given by

**FIG. 1.**
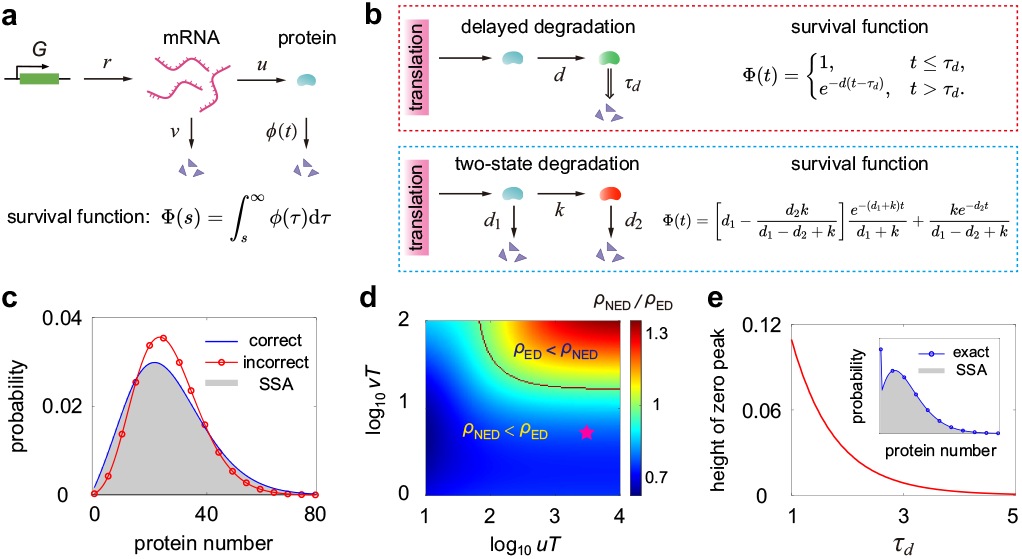
Two-stage model. **(a)** Schematic of the model, which involves transcription, translation, mRNA decay, and protein decay. **(b)** Two protein degradation mechanisms and their survival functions — the delayed degradation [5–8] and two-state degradation [3, 4, 14]. **(c)** Comparison between the correct solution given in Eq. (11) (blue curve), the incorrect solution given in [8] (red circles), and the numerical solution obtained from the SSA (gray region) for the two-stage model with delayed protein degradation. **(d)** Heat plot of *ρ*NED*/ρ*ED as a function of *uT* and *vT*. The red star shows the medians of *vT* and *uT* in mouse fibroblasts [3, 15]. **(e)** Height of the zero peak of the protein distribution versus *τd* in the case of deterministic protein degradation. See [16] for the technical details of the figure.

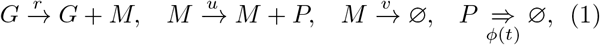

where *r* is the transcription rate, *u* is the translation rate, and *v* is the mRNA decay rate. The last reaction describes protein decay with *ϕ*(*t*) being the probability density of the protein degradation time. For exponentially degraded (ED) proteins, the above model reduces to the conventional two-stage gene expression model [13].

Since protein degradation is non-exponential, the underlying stochastic dynamics is non-Markovian. We next develop an effective Markovian approach to find the exact solution of the non-Markovian system. Let *t* > 0 be a fixed time point, let *M* (*t*) and *N* (*t*) denote the copy numbers of mRNA and protein at time *t*, respectively, and let *p*_*m*,*n*_(*t*) = ℙ (*M* (*t*) = *m, N* (*t*) = *n*) denote the probability of having *m* copies of mRNA and *n* copies of protein at time *t*. For each *s* ≤ *t*, let *N* (*s*; *t*) denote the number of protein molecules that are present at time *s* and survive (i.e. are not degraded) until time *t*. Clearly, we have *N* (*s*; *t*) ≤ *N* (*s*) for each *s* ≤ *t* and *N* (*t*; *t*) = *N* (*t*).

We now examine the stochastic dynamics of the binary process (*M* (*s*), *N* (*s*; *t*)), *s* ≤ *t*. We emphasize that while the original process (*M* (*t*), *N* (*t*)) is non-Markovian, the effective process (*M* (*s*), *N* (*s*; *t*)) becomes Markovian. We claim that at each time *s* ≤ *t*, the effective process can only make the following three types of transitions:

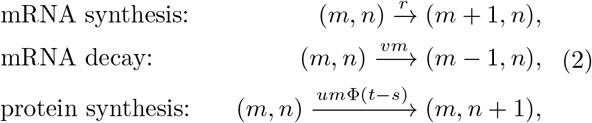

where 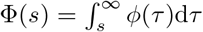 is the survival function of the protein degradation time distribution.

We next prove this fact. Since *M* (*s*) is Markovian, the synthesis and decay of mRNA will result in transitions from (*m, n*) to (*m* + 1, *n*) and (*m* − 1, *n*), respectively. To understand how *N* (*s*; *t*) evolves, note that *N* (*s*; *t*) must increase with *s*, which means that *N* (*s*; *t*) ≤ *N* (*s*^′^; *t*) for any *s < s*^′^ ≤ *t*. This is because the protein molecules that exist at *s*^′^ and survive until time *t* include (i) those molecules that already exist at time *s* and survive until time *t* and (ii) those molecules produced during the time interval (*s, s*^′^] and surviving until time *t*. Since *N* (*s*; *t*) increases with *s*, the transition from (*m, n*) to (*m, n* + 1) is allowed for the new process; however, the transition from (*m, n*) to (*m, n* − 1) is forbidden. At each time *s* ≤ *t*, the transition rate from (*m, n*) to (*m, n* + 1) should be the product of the rate *um* at which protein is produced and the probability Φ(*t* − *s*) that a newly produced protein molecule survives until time *t*.

Let *p*_*m*,*n*_(*s*; *t*) = ℙ (*M* (*s*) = *m, N* (*s*; *t*) = *n*). Since the effective process (*M* (*s*), *N* (*s*; *t*)) is Markovian and its all transitions are given in Eq. (2), the evolution of *p*_*m*,*n*_(*s*; *t*) is governed by the chemical master equation (CME)

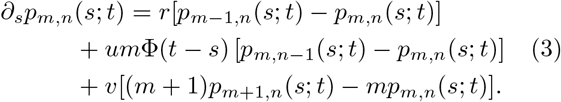

Clearly, the effective process (*M* (*s*), *N* (*s*; *t*)) governs the time evolution of the effective Markovian reaction system

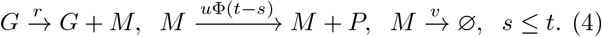

Note that there are two crucial characteristics for the effective Markovian dynamics (EMD): (i) the EMD does not include protein decay; (ii) for each fixed time *t* > 0, the EMD is a finite-time dynamics until time *t* and its translation rate also depends on the terminal time *t*.

Since *N* (*t*; *t*) = *N* (*t*), the original non-Markovian dynamics and the EMD must have the same distribution of mRNA and protein numbers at the terminal time *t*, which means that *p*_*m*,*n*_(*t*; *t*) = *p*_*m*,*n*_(*t*). Hence if the EMD can be solved exactly in time, then we automatically obtain the analytical solution of the non-Markovian system. To solve the EMD, we define the generating functions

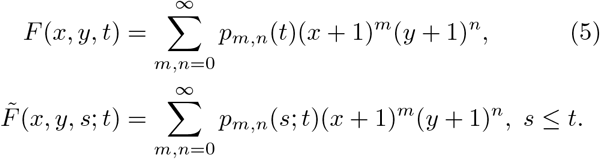

Clearly, we have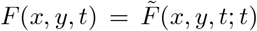. Using these generating functions, Eq. (3) can be converted into the first-order linear partial differential equation (PDE)

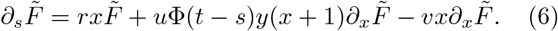

Solving 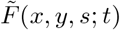 from Eq. (6) and then taking *s* = *t* gives the explicit expression of *F* (*x, y, t*). Throughout this paper, we assume that initially there are no mRNA and protein molecules in the cell, i.e. *M*_0_ = *N*_0_ = 0. In [16], we show that under this initial condition, the generating function can be obtained in closed form as

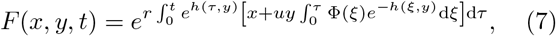

Where 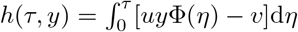. The time-dependent joint distribution *p*_*m*,*n*_(*t*) of mRNA and protein numbers can be recovered from *F* (*x, y, t*) as

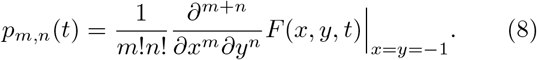

Moreover, taking *t* → ∞ in Eq. (7), we obtain the generating function of the steady-state joint distribution

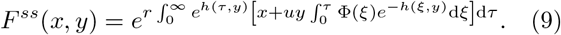

We then use Eq. (9) to derive the exact joint distribution of mRNA and protein copy numbers in the case of delayed protein degradation (Fig. 1(b), upper panel). Following [5–8], we model protein decay as a reaction that is first initiated at rate *d* and complete after a deterministic time *τ*_*d*_ after initiation. The fixed time delay is valid when the degradation process is tightly regulated or when it is composed of many steps [6]. In this case, the survival function Φ(*t*) is given by

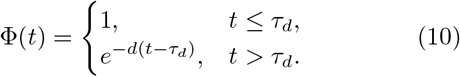

Inserting Eq. (10) into Eq. (9) yields

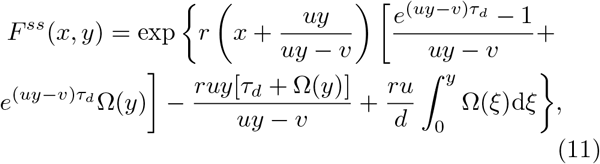

where Ω(*y*) = _1_*F*_1_(1; 1 + *v/d*; *uy/d*)*/v* with _1_*F*_1_ being the confluent hypergeometric function. When *τ*_*d*_ = 0, the above solution reduces to the classical result for exponential protein degradation [17].

We emphasize that the marginal protein distribution for the same model has already been solved in [8], but that solution is incorrect. In [16], we briefly explain how the incorrect solution comes from. Fig. 1(c) compares the true exact solution given in Eq. (11), the false one given in [8], and the numerical one obtained using the stochastic simulation algorithm (SSA). In [16], we also show that the exact solutions obtained in [13, 17, 18] for simpler models can be viewed as special cases of our solution.

### Two-state degradation

A recent study measured the decay profiles for thousands of proteins and found that many proteins are degraded non-exponentially [3]. The survival functions Φ(*t*) of those NED proteins are all well fitted to a two-state degradation mechanism (Fig. 1(b), lower panel) [3, 14]. Here newly synthesized proteins first populate state *P*_1_, from which they can either be degraded with rate *d*_1_ or convert to another state *P*_2_ with a lower decay rate *d*_2_ *< d*_1_. Moreover, it has been shown [3] that the median lifetimes for ED and NED proteins in mouse fibroblasts are similar (≈ 52 h; see [16] for the detailed estimation); however, the correlation coefficient between mRNA and protein numbers for NED proteins, *ρ*_NED_, is on average less than that for ED proteins, *ρ*_ED_ — the median of *ρ*_NED_*/ρ*_ED_ is only 0.76 (see Fig. S2 in [19]).

We now use our analytical results to explain this phenomenon. Recall that for ED proteins, the correlation between mRNA and protein numbers is given by [20]

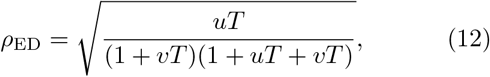

where *T* is the mean protein lifetime, *vT* is the ratio of the mean lifetimes of protein and mRNA, and *uT* is ratio of the mean numbers of protein and mRNA in a single cell. In [3, 14], the gene expression dynamics for NED proteins is modeled by the following two-stage model with two-state protein degradation:

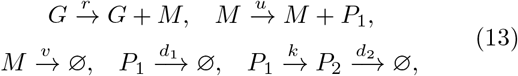

where *d*_*i*_, *i* = 1, 2 is the degradation rate of *P*_*i*_ and *k* is the conversion rate from *P*_1_ to *P*_2_. If we do not distinguish between *P*_1_ and *P*_2_ and focus on the total number of them, then the above model is equivalent to the non-Markovian model given in Eq. (1) with decay time distribution

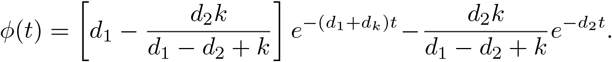

The mRNA-protein correlation for NED proteins is complicated and can be found in [16]. The medians of *uT*, *vT*, *d*_1_, *d*_2_, and *k* in mouse fibroblasts can be determined using experimental data in [3, 15] and the specific values can be found in Table I. For example, the median mRNA number in mouse fibroblasts in 17 and the median protein number is 50000 [15]. Hence *uT* is estimated to be 50000*/*17 ≈ 2900.

Fig. 1(d) shows the heat plot of *ρ*_NED_*/ρ*_ED_ as a function of *uT* and *vT*, where the mean lifetimes for ED and NED proteins are assumed to be equal. Interestingly, we find that *ρ*_NED_ *< ρ*_ED_ whenever *vT* is relatively small. This is fully consistent with our theoretical results. In [16], we prove that *ρ*_NED_ is less than *ρ*_ED_ whenever

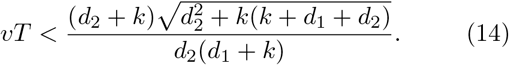

From Table I, the median of the right-hand side of Eq. (14) in mouse fibroblasts is estimated to be 15.79, while the median of *vT* is only 5. This explains why NED proteins on average have a smaller mRNA-protein correlation coefficient than ED proteins. More importantly, if we choose *vT* and *uT* to be their medians in Table I, then we have *ρ*_NED_*/ρ*_ED_ ≈ 0.81 (Fig. 1(d), red star). This is close to the value of 0.76 observed in experiments [19].

**TABLE 1.** Medians of *uT*, *vT*, *d*1, *d*2, and *k* in mouse fibroblasts (NIH 3T3). See [16] for the detailed estimation of these parameters.

| | $uT$ | $vT$ | $d_1$ (h<sup>-1</sup>) | $d_2$ (h<sup>-1</sup>) | $k$ (h<sup>-1</sup>) |
| --- | --- | --- | --- | --- | --- |
| median | 2900 [15] | 5 [15] | 0.427 [3] | 0.0087 [3] | 0.224 [3] |

### Three-stage model

We next apply our EMD method to a three-stage gene expression model involving gene state switching, transcription, and translation (Fig. 2(a)):

**FIG. 2.**
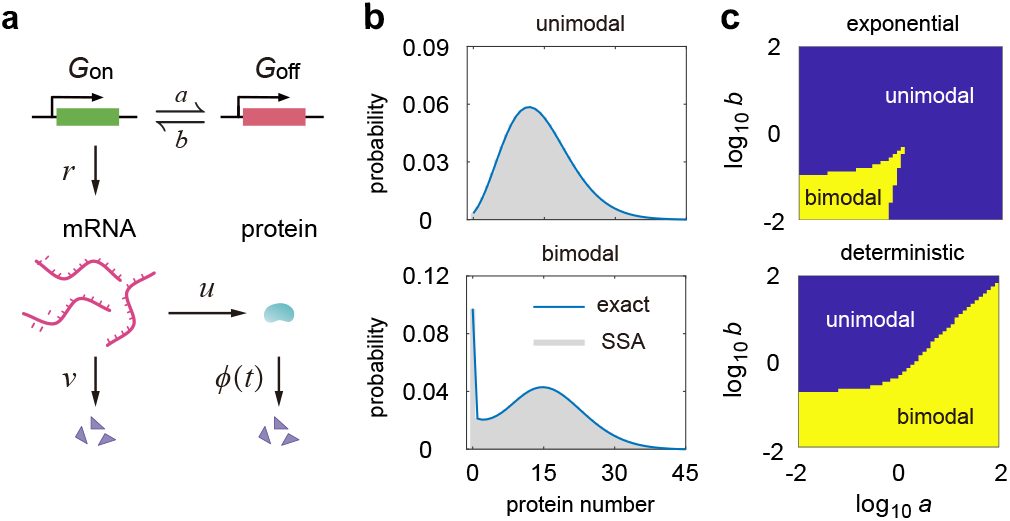
Three-stage model. **(a)** Schematic of the model, which involves transcription, translation, mRNA decay, protein decay, and gene state switching. **(b)** Comparison of the analytical protein distribution given in Eq. (18) (blue curve) with the SSA (gray region) for the three-stage model with deterministic protein decay. **(c)** Phase diagrams in the *a*-*b* plane for the three-stage model with exponential (upper) and deterministic (lower) protein decay. See [16] for the technical details of the figure.

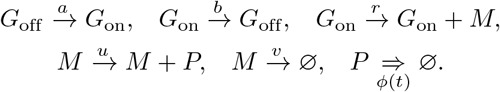

Here *a* and *b* are the switching rates between the active state *G*_on_ and inactive state *G*_off_ of the gene, *r* is the transcription rate, *u* is the translation rate, *v* is the mRNA decay rate, and *ϕ*(*t*) is the decay time distribution of each protein molecule. Transcription can only occur when the gene is in the active state. We now use the EMD approach to solve this non-Markovian model. Let *t* > 0 be a fixed time point, let *I*(*t*) denote the state of the gene at time *t*, let *p*_*m*,*n*_(*t*) = ℙ (*M* (*t*) = *m, N* (*t*) = *n*) denote the probability of having *m* copies of mRNA and *n* copies of protein, and let *F* (*x, y, t*) denote its generating function.

We next examine the dynamics of the effective ternary process (*I*(*s*), *M* (*s*), *N* (*s*; *t*)), *s* ≤ *t*, where again, *N* (*s, t*) denotes the number of protein copies that are present at time *s* and survive until time *t*. Similarly to the two-stage model, the effective process governs the time evolution of the effective Markovian reaction system

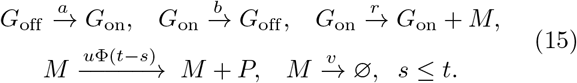

To proceed, let

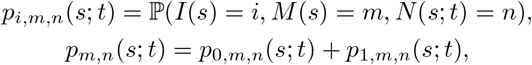

where *i* = 0, 1 represent the inactive and active gene states, respectively, and *p*_*m*,*n*_(*s*; *t*) denotes the joint distribution of mRNA and protein numbers for the EMD. To solve the EMD, we define the generating functions

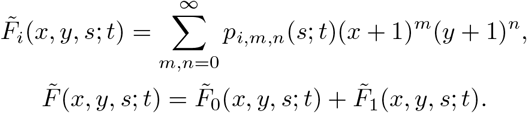

Again, we have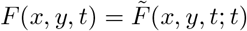. Similarly to the two-stage model, the generating functions for the EMD satisfy the first-order linear PDEs [16]

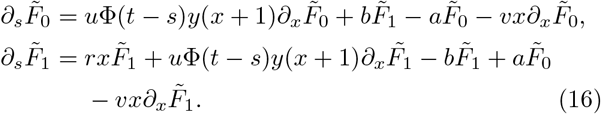

In general, these PDEs are not solvable. However, it can be solved exactly in some special cases.

We next consider the case where protein decay requires a deterministic time *τ*_*d*_ to complete. Deterministic gene product degradation is also studied extensively in the literature [9–11]. Under this assumption, the survival function Φ(*t*) is given by

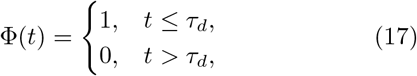

and clearly, it is piecewise constant. Note that if Φ(*t*) is constant, then the EMD given in Eq. (15) is a classical three-stage gene expression model without protein decay, and its exact solution has been obtained in [21]. Now since Φ(*t*) is piecewise constant, the EMD is still solvable.

Specifically, solving 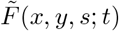 from Eq. (16) and then taking *s* = *t* gives the exact solution of *F* (*x, y, t*). Finally, taking *t* → ∞ in *F* (*x, y, t*), we obtain the generating function *F* ^*ss*^(*x, y*) of the steady-state joint distribution of mRNA and protein numbers [16]:

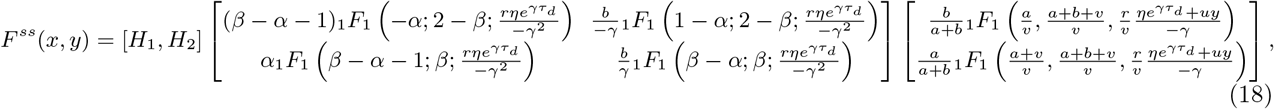

where

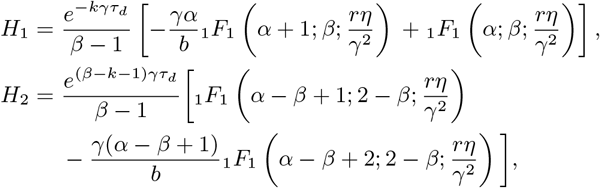

and the constants *k, α, β, γ, η* are given by

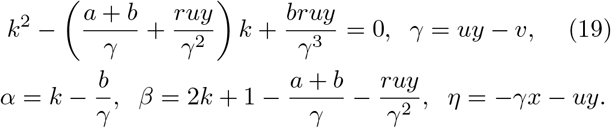

Fig. 2(b) compares the analytical protein number distribution with the SSA. As expected, the two solutions agree perfectly. According to simulations, the protein distribution can be either unimodal or bimodal. To understand how delayed protein degradation affects bimodality, we illustrate the *a*-*b* phase diagrams for the three-stage model with exponential and delayed protein degradation in Fig. 2(c). For exponential degradation, bimodality only occurs when the gene switches slowly between the active and inactive states [22]. Interestingly, the bimodal parameter region becomes significantly larger for delayed degradation. In the End Matter, we use our analytical results to explain why bimodality is greatly enhanced in the presence of delayed protein degradation.

### Multistate model

Our EMD approach can be further applied to obtain analytical results for multistate gene expression models. To see this, we consider the following multistate model with deterministic protein decay:

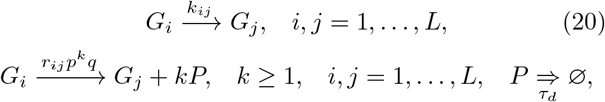

where *q* = 1 − *p*. Here the gene of interest can switch freely between *L* states *G*_1_, …, *G*_*L*_ with rates *k*_*ij*_. These gene states correspond to different conformational states during chromatin remodeling or different binding states with transcription factors and RNA polymerases [23]. In each gene state *G*_*i*_, protein is produced in a bursty manner with frequency *r*_*ij*_ and random size *k* having the geometric distribution *p*^*k*^*q*; following [10], after protein synthesis, we allow the change of gene state from *G*_*i*_ to *G*_*j*_.

Note that this multistate model has also been used to describe the dynamics of nascent RNA [9, 10]. Real-time observation of transcription *in vivo* has shown that while transcription initiation is a stochastic process, elongation and termination are fairly deterministic [12]. If the molecule *P* is understood to be nascent RNA, then the reaction *P* ⇒ ∅ means that a nascent transcript becomes a mature transcript after a fixed time *τ*_*d*_.

We stress that in previous studies [9, 10], the multistate model with deterministic time delay is only analytically solved when (i) the protein is produced in a non-bursty manner and (ii) there is only one active gene state and the other gene states are all inactive. The exact solution when any one of the two restrictions is violated is still an open problem [11].

Interestingly, the EMD approach can be applied to derive the analytical protein distribution for the reaction scheme given in Eq. (20) when the above two restrictions are broken. The idea is to convert the non-Markovian dynamics into the following EMD:

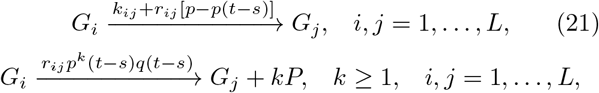

where *p*(*t*) = *p*Φ(*t*)*/*[*p*Φ(*t*) + *q*], *q*(*t*) = 1 − *p*(*t*), and the survival function Φ(*t*) is given in Eq. (17). Again, the non-Markovian dynamics and the EMD have the same protein distribution at the terminal time *t*. The exact solution for this model and its detailed derivation can be found in the End Matter.

## Conclusions

In this study, we developed an EMD method of solving a large class of non-Markovian gene expression models with complex protein degradation mechanisms. The central idea is to convert non-Markovian dynamics into EMD so that they share the same mRNA and protein number distributions at any fixed time. Then non-Markovian systems can be solved in closed form whenever the EMD has an exact solution. Our method was then applied to obtain analytical results for some important models that fail to be solved using conventional queueing theory. Our results also provide a theoretical explanation of why NED proteins on average have smaller mRNA-protein correlation than ED proteins.

## Acknowledgements

Y. L. acknowledges support from NSFC grant No. 12401629. C. J. acknowledges support from NSFC grants No. 12271020 and No. U2230402.

## Appendix A Bimodality in the presence of delayed protein degradation

According to simulations, our model can produce a unimodal or bimodal protein number distribution (Fig. 2(b)). The bimodal one is of particular interest because it indicates the separation of isogenic cells into two distinct phenotypes. To understand how delayed protein degradation affects bimodality, we illustrate the *a*-*b* phase diagrams for the three-stage model with exponential and delayed (deterministic) protein degradation in Fig. 2(c). For exponential degradation, bimodality only occurs when the gene switches slowly between the active and inactive states [22]. Interestingly, the bimodal parameter region becomes significantly larger for delayed degradation — bimodality takes place when the gene activation rate is much larger than the gene inactivation rate (*a* ≫ *b*).

Note that when *a* ≫ *b*, the gene is mostly active and hence the three-stage model reduces to the two-stage model shown in Fig. 1(a). Taking *d* → ∞ in Eq. (11) and then setting *x* = 0, we obtain the generating function of the steady-state protein number distribution for the two-stage model with deterministic protein decay:

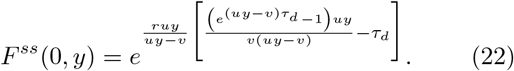

In most cell types such as bacteria, yeast, and mammalian cells, mRNA often decays much faster compared to its protein counterpart [24]. Note that in the presence of deterministic protein decay, *vτ*_*d*_ represents the ratio of the mean protein lifetime to mean mRNA lifetime. In mouse fibroblasts (NIH 3T3), mRNA and protein have median half-lives of 9 h and 46 h, respectively [15]; in this case, we have *vτ*_*d*_ ≈ 5. The value of *vτ*_*d*_ is even larger in bacteria and yeast [24].

When *vτ*_*d*_ ≫ 1 and *u/v* is strictly positive and bounded, it is well-known that protein synthesis will occur in a bursty manner with the burst size having a geometric distribution [13, 25]. In this case, the two-stage model further reduces to the following bursty model:

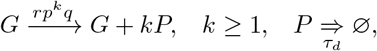

where *p* = *u/*(*u* + *v*) and *q* = 1 − *p*. Note that in this model, protein synthesis occurs in bursts with frequency *r* and random size *k* sampled from a geometric distribution with parameter *p*, in agreement with experiments [26].

Taking the limit of *vτ*_*d*_ → ∞ while keeping *B* = *u/v* as constant in Eq. (22), we obtain the steady-state generating function for the above bursty model:

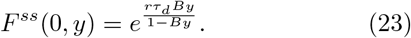

Finally, the steady-state protein number distribution can be recovered by taking the derivatives of the generating function at *y* = −1 and is given by

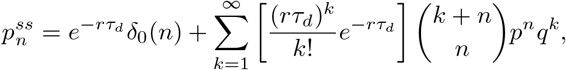

where *δ*_0_(*n*) is Kronecker’s delta that equals 1 when *n* = 0 and equals 0 otherwise. Clearly, the protein distribution is a mixture of many negative binomial distributions with the weights having a Poisson distribution. The first component in the mixture produces the peak at zero with height 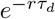, and the remaining components produce the non-zero peak. This reveals the origin of bimodality in the presence of delayed protein decay. Our theory predicts that the zero peak becomes higher as *τ*_*d*_ decreases. This is consistent with our simulations in Fig. 1(e).

## Appendix B: Analytical solution for the multistate model

Here we derive the analytical solution for the following multistate gene expression model with deterministic protein degradation:

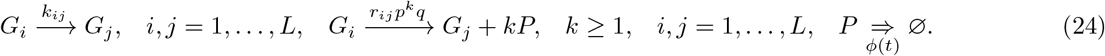

where *ϕ*(*t*) = *δ*(*t* − *τ*_*d*_) with *δ*(*t*) being the Dirac Delta function. Let *p*_*i*,*n*_(*t*) = ℙ (*I*(*t*) = *i, N* (*t*) = *n*) denote the probability of having *n* copies of protein when the gene is in state *i*. We next examine the dynamics of the effective process (*I*(*s*), *N* (*s*; *t*)), *s* ≤ *t*, where again, *N* (*s*; *t*) denotes the number of protein copies that are present at time *s* and survive until time *t*.

Since *N* (*s*; *t*) increases with *s*, the transitions from (*i, n*) to (*i, n* − 1) are forbidden for the effective process. Note that for the effective process, the transitions from (*i, n*) to (*j, n*) can be achieved either via the reaction *G*_*i*_ → *G*_*j*_ or via the reactions *G*_*i*_ → *G*_*j*_ + *kP*, *k* ≥ 1 with all the protein molecules produced in a single burst being degraded before time *t*. Hence the transition rate from (*i, n*) to (*j, n*) for the effective process is given by

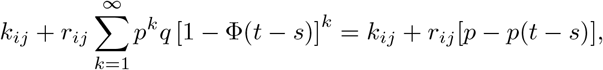

where *p*(*t*) = *p*Φ(*t*)*/*[*p*Φ(*t*) + *q*]. Similarly, the transitions from (*i, n*) to (*j, n* + *k*) for the effective process can be achieved via the reactions *G*_*i*_ → *G*_*j*_ + *lP*, *l* ≥ *k* with *l* − *k* protein molecules being degraded before time *t*. Hence the transition rate from (*i, n*) to (*j, n* + *k*) for the effective process is given by

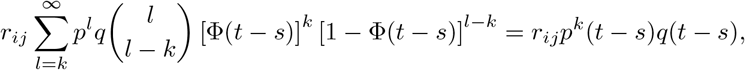

where *q*(*t*) = 1 − *p*(*t*). In summary, we have proved that the EMD for the multistate model is governed by Eq. (21) in the main text.

We next solve the EMD for the multistate model with deterministic protein decay. In this case, the survival function is given by Eq. (17). When *t* ≤ *τ*_*d*_, we have Φ(*t* − *s*) = 1 and hence the EMD is governed by the reactions

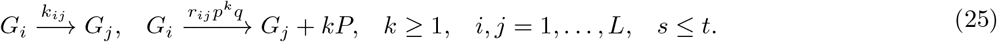

The evolution of *p*_*i*,*n*_(*s*; *t*) is then governed by the CMEs

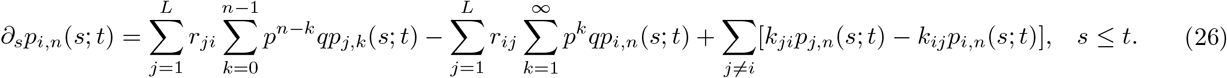

To solve the CMEs, we define the generating functions:

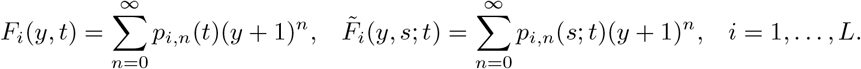

Using the generating functions, Eq. (26) can be converted into the system of ordinary differential equations (ODEs)

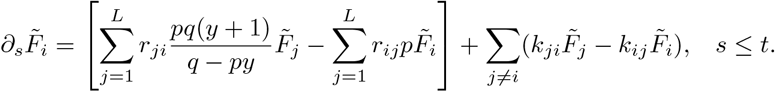

Note that the right-hand side of the above system of ODEs is independent of the time variable *s*. Hence it can be viewed as a linear system of ODEs with constant coefficients and its analytical solution is given by

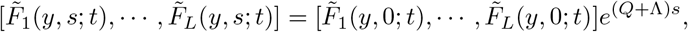

where the matrix *Q* = (*q*_*ij*_)_1≤*i*,*j*≤*L*_ is defined by

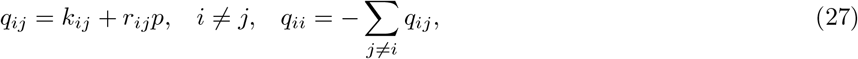

the matrix Λ = (*λ*_*ij*_)_1≤*i*,*j*≤*L*_ is defined by *λ*_*ij*_ = *r*_*ij*_*py/*[*q* − *py*], and 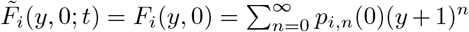 are the generating functions associated with the initial conditions. Note that *q*_*ij*_ characterizes both free gene state switching via the reaction *G*_*i*_ → *G*_*j*_ and gene state switching induced by protein synthesis via the reaction *G*_*i*_ → *G*_*j*_ + *kP*. Since the non-Markovian model and the EMD have the same protein number distribution at the terminal time *t*, we must have 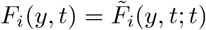. This shows that

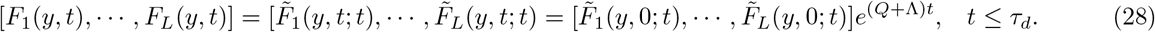

This gives the analytical time-dependent generating function when *t* ≤ *τ*_*d*_.

When *t* ≥ *τ*_*d*_, we have Φ(*t* − *s*) = 0 when *s* ≤ *t* − *τ*_*d*_ and Φ(*t* − *s*) = 1 when *t* − *τ*_*d*_ *< s* ≤ *t*. We next focus on these two cases separately. When *s* ≤ *t* − *τ*_*d*_, the EMD is governed by the reactions

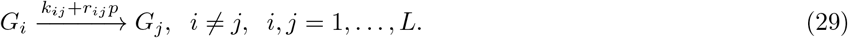

When *t* − *τ*_*d*_ *< s* ≤ *t*, the EMD is still governed by the reactions given in Eq. (25). Straightforward computations show that the exact solution of the EMD is given by [16]

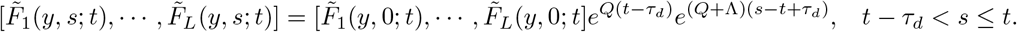

Since the EMD and the non-Markovian model have the same protein number distribution at the terminal time *t*, we must have 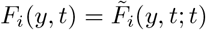. This indicates that

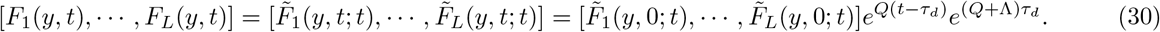

Eqs. (28) and (30) give the complete time-dependent generating function for the multistate model with deterministic protein degradation.

Finally, we focus on the steady-state protein distribution. Note that the matrix *Q* defined in Eq. (27) is the generator matrix of an *L*-state Markovian system. Without loss of generality, we assume that *Q* is irreducible. To proceed, we make a crucial observation that

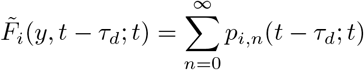

is actually the probability of the gene being in state *G*_*i*_ at time *t* − *τ*_*d*_ since protein molecules surviving at *t* can only be produced at time *s* satisfying *t* − *τ*_*d*_ *< s* ≤ *t*. As *t* → ∞, the *L*-state Markovian system will reach a steady state, which implies that

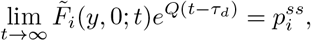

where 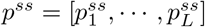 is the steady-state distribution of the *L*-state Markovian system satisfying

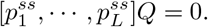

Take *t* → ∞ in Eq. (30), we obtain the steady-state generating function of the protein number distribution:

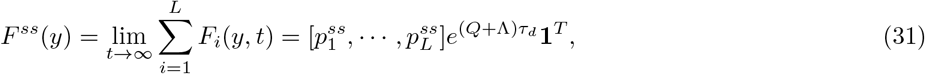

where **1** = [1, …, 1] is the row vector whose components are all 1. In [16], we also show that the exact solutions obtained in [9, 10] for simpler models can be viewed as special cases of our solution. Moreover, we emphasize that even when there is only one active gene state and the other gene states are all inactive, our solution still has a much more concise form than the one derived in [10].

